# PROSTATA: Protein Stability Assessment using Transformers

**DOI:** 10.1101/2022.12.25.521875

**Authors:** Dmitriy Umerenkov, Tatiana I. Shashkova, Pavel V. Strashnov, Fedor Nikolaev, Maria Sindeeva, Nikita V. Ivanisenko, Olga L. Kardymon

**Affiliations:** Sber AI Lab, Moscow, Russia; AIRI, Moscow, Russia; Laboratory of Computational Proteomics, Institute of Cytology and Genetics SB RAS, Novosibirsk, Russia

## Abstract

Accurate prediction of change in protein stability due to point mutations is an attractive goal that remains unachieved. Despite the high interest in this area, little consideration has been given to the transformer architecture, which is dominant in many fields of machine learning. In this work, we introduce PROSTATA, a predictive model built in knowledge transfer fashion on a new curated dataset. PROSTATA demonstrates superiority over existing solutions based on neural networks. We show that the large margin of improvement is due to both the architecture of the model and the quality of the new training data set. This work opens up opportunities for developing new lightweight and accurate models for protein stability assessment. PROSTATA is available at https://github.com/AIRI-Institute/PROSTATA.

## 1 Introduction

Quantitative prediction of the effects of single amino acid substitutions on protein stability is a major problem that remains unresolved. Protein stability is related to its structure, function, and molecular evolution. The prediction of protein stability is part of a broader issue of predicting evolutionary fitness and the phenotypic effects of genomic variations.

Accurate predictions of changes in protein stability caused by mutations provide crucial insight into how proteins fold and function, and also have important applications in the bioindustry. Indeed, an exponential increase of chemical reaction rates with temperature makes an improvement in molecular stability a desirable outcome.

Change in protein stability due to mutation is usually expressed through either (un)folding free energy ΔΔ*G* or the melting temperature *T*_*m*_. ΔΔ*G* is often referred to as thermodynamic stability. It is determined from the thermodynamic cycle as the difference between the changes in free energy upon folding of the wild-type protein and its mutated form. Since free energy depends on temperature, the standard reference temperature of 298 K (room temperature) is universally agreed upon and is used by default. *T*_*m*_ is the thermal denaturation midpoint, that is, the temperature at which the concentrations of folded and denatured forms of the protein are equal. The difference between *T*_*m*_s of mutated and wild-type proteins is a measure of thermal stability 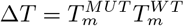.

Thermodynamic stability ΔΔ*G* and thermal stability Δ*T*_*m*_ are often used as proxies. Both of these descriptors are protein dependent and show linear anticorrelation. However, the correlation between ΔΔ*G* and Δ*T*_*m*_ differs from -1 because ΔΔ*G* is often considered at standard temperature, not at 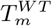. This implies that ΔΔ*G* and Δ*T*_*m*_ cannot be used interchangeably as a measure of protein stability and should be considered independent (Pucci et al., 2018).

The experimental measurement of ΔΔ*G* and Δ*T*_*m*_ is a difficult and non-trivial task. Different experimental techniques, protocols, and conditions often produce heterogeneous or even contradicting results (Wright et al., 2017). Moreover, experimental results naturally biased towards destabilizing mutations due to the fact that wild-type sequences are, in a sense, locally optimal in terms of stability. Thus, the scarcity and misalignment of the experimental data lead to imbalances in the available datasets for training protein stability prediction models (Louis and Abriata, 2021; Bæk and Kepp, 2022).

Machine learning has irreversibly changed the landscape of computational biology and molecular modeling over the last years. A plethora of tools designed to predict protein stability perfectly illustrate this change (Horne and Shukla, 2022). We can roughly divide all the tools into three categories: (a) structural modeling methods employing some kind of empirical energy function, (b) ‘simple’ machine learning tools based on such methods as support vector machines (SVM), and (c) deep neural networks, mostly convolutional neural networks (CNNs).

First category includes classical methods like Rosetta (Leman et al., 2020), as well as new developed methods like PoPMuSiC (Dehouck et al., 2011). Rosetta is a suite of macromolecular modeling programs (Leman et al., 2020).

Rosetta generates and refines 3D structural models of the mutated protein and its corresponding wild-type structure and then calculates the energy difference between them. Rosetta employs an energy function in the form of a linear combination of physics-based and knowledge-based contributions. PoPMuSiC is a knowledge-based predictor using a statistical energy function trained on a large experimental data set (Dehouck et al., 2011).

Classic machine learning models are by far the most populous category of tools for predicting protein stability (Horne and Shukla, 2022). For example, DDGun (Montanucci et al., 2022) is an untrained method that combines 3 evolutionary sequence-based scores in a linear combination. Its structure-based version, DDGun3D, in addition to the 3 scores used in DDGun introduces another term calculated through a statistical potential. K. Bæk and K. Kepp introduced simple interpretable linear regression models that achieve accuracy similar to that of more complex prediction methods (Caldararu et al., 2021; Bæk and Kepp, 2022). These regression models use only three descriptors, namely, relative solvent accessibility, volume difference, and hydrophobicity difference.

Recently, methods from the last category based on deep learning approaches became popular. In particular, this category includes such methods as DeepDDG (Cao et al., 2019), ThermoNet (Li et al., 2020), SCONES (Samaga et al., 2021), ACDC-NN (Benevenuta et al., 2021), ProS-GNN (Wang et al., 2021). Despite the more complex model architecture, this class of methods still does not have a clear advantage over others (Pucci et al., 2018; Pancotti et al., 2022).

The performance of the model depends on both the architecture and the training set. Most of the datasets are derived from the ProTherm database (Nikam et al., 2020), the largest collection of experimental mutation data. However, actual datasets for model training and testing could be combined in different ways according to (1) experimental conditions (e.g. pH, temperature), (2) symmetries of stabilizing and destabilizing mutations, and (3) proteins sequence similarities.

The question of how to aggregate data across experimental conditions is still open, while the importance of training set symmetry has been shown in work by Pucci et al. (Pucci et al., 2018). The authors presented symmetric test set called Ssym to compare the performance of various models in stabilizing and destabilizing mutations. The results show that most models learned on non-symmetric training set shift their predictions towards destabilizing mutations.

For a correct comparison of the developed models, it is important that the test set does not overlap with the training set. In work by Li B. et al., it is shown that most of the data sets used to train models contain proteins from the Ssym data set (Li et al., 2020). This fact makes a fair comparison unfeasible. To address this issue, Pancotti et al. developed a new test set called S669 that does not overlap with the widely used training sets (Pancotti et al., 2022).

In summary, multiple approaches have been developed to protein stability prediction, however, improving the accuracy of prediction is still of great importance. At the same time, transformers, widely used in many areas of AI since their discovery by Vaswani et al. in 2017 (Vaswani et al., 2017), have not yet found their way into considered field. In this work, we show that the transformer architecture can be successfully applied to predict relative protein stability. In addition, we built a new data set, merging data according to best practice from previously works and ambiguities were manually curated.

## 2 Materials and methods

### 2.1 External datasets

In this work, to compare our model with other NN methods, we used the original training datasets for the corresponding models where such data were readily available in unified format. We used Q3421 from STRUM (Quan et al., 2016), Q3488 from ThermoNet (Li et al., 2020), the widely used S2648 training set provided by Dehouck Y. *et al*. (Dehouck et al., 2011), and additional data from Varibench for ACDC-NN models (Benevenuta et al., 2021). Data sets Q3488 and Q3421 were used to assess the effect of a non-symmetric training set on PROSTATA prediction.

The commonly used test sets Ssym (Pucci et al., 2018) and S669 (Pancotti et al., 2022) were chosen as test sets to evaluate the models.

### 2.2 Dataset construction

We constructed our own data set based on relevant sets from the VariBench portal (Nair and Vihinen, 2013), including popular training sets such as PoPMuSic-2.0 (S2648) (Dehouck et al., 2011), ThermoNet (Q3214) (Li et al., 2020) and VariBench (Nair and Vihinen, 2013) (Supplementary Table S1). Data were merged and hand-checked.

Since our model does not use experimental conditions as features, we have aggregated samples by a combination of PDB ID, PDB chain, and mutation code (position and residues in it before and after mutation), hereinafter referred to as ID. Data were averaged over experimental pH and temperature (T) and pooled in five steps.

1. **Split the data**. All samples were divided into two groups according to whether *pH* and *T* were available (Group I) or not (Group II).
2. **Select core samples**. The samples in Group I with *pH* and *T* closest to the standard values (*pH* = 7 and *T* = 25^*°*^*C*) were selected by ID.
3. **Select additional samples**. From the remaining samples of Group I, for each core sample, we selected the corresponding samples with *pH* = *pH*_*core*_ ± 0.5 and *T* = *T*_*core*_ ± 10^*°*^*C*. Samples with IDs unique to Group II were also selected.
4. **Average** ΔΔ*G* **over mutations**. For each ID of the selected samples, the ΔΔ*G* values were calculated as the mean of the experimental ΔΔ*G* values.
5. **Discard inconsistencies**. To construct the final dataset, the samples with conflicting ΔΔ*G* values (e.g. the opposite signs of ΔΔ*G* values or variance of ΔΔ*G* greater than 5 kJ/mol) were filtered out.

The final dataset contains 5,196 samples (Supplementary Table S2). To make a fair comparison with other methods, we collected the corresponding protein sequences for our dataset and the test sets (Ssym and S669) to filter out similar sequences between them. Sequences were clustered with 75% identity using the MMSeqs software (Steinegger and Söding, 2017). Samples from our dataset not assigned to any clusters in the test sets were included in a training set. To avoid an imbalance of the data set in favor of destabilizing mutations, for each mutation in the training set, a reversed mutation was also included (Supplementary Table S3, S4).

To provide most accurate results to the end users, the publicly available final version of the model was trained using the whole dataset without filtering out any sequences. The whole dataset comprises our training dataset combined with test sets and contains 10,542 amino acids substitutions (Supplementary Table S5).

### 2.3 Model architecture

We treat the prediction of the mutation effect on protein thermostability as a regression task for two sequences, the wild-type and mutated. Using transformer models for this task is a two-step process. First, a model pre-trained on a large corpus of unlabelled data is used to extract the representations of the sequences. Second, the sequence representations of wild-type and mutated proteins are combined into a single representation that is used to predict the target value. Our models consist of a transformer backbone that produces the embeddings for wild-type and mutated proteins and the regression head that combines the embeddings in various ways to predict ΔΔ*G* (Figs. 1–3). The final predictions are made by an ensemble of five models.

**Fig. 1.**
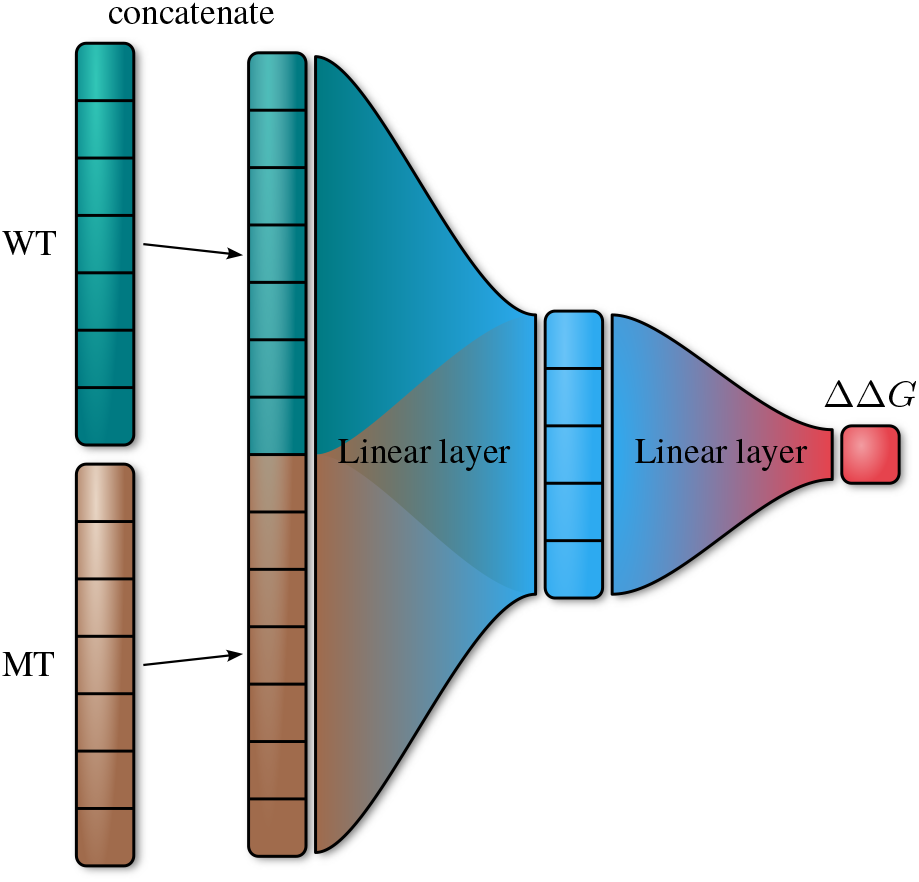
Architecture of the model that uses the concatenation of token embeddings in mutation position of wild-type (WT) and mutated (MT) protein passed as input to the neural network with one hidden layer.

#### 2.3.1 Sequence embedding with transformer backbone

There are several transformer models pre-trained on unlabelled sequential protein data, such as ProtTrans, ProteinBERT, ESM, ESM-2 (Lin et al., 2022). For this work, we settled on using one of the ESM-2 models as the embedding backbone. The ESM-2 is a family of models of different sizes with parameter count ranging from 8 million to 15 billion, with larger models producing better protein representations. For this work, we employ the ESM-2 model with 650 million parameters, as it is the largest model that can be trained on a 32 GB GPU. This model produces embeddings of size 1280 per residue. Larger models could potentially be used to achieve higher quality at the expense of much longer training and inference times. During sequence embedding, the model calculates representations for each amino acid in the sequence. Additionally, the model calculates representations for special tokens, namely, the CLS token that is inserted at the beginning of each sequence and the END token that is appended to each sequence. The output of the transformer backbone for each protein sequence of length *N* is a vector of size (*N* + 2) *×* 1280.

#### 2.3.2 Regression head

The second step in the regression pipeline is to combine wild-type (WT) and mutated (MT) embeddings into a joint representation that is used as input for a linear regression head. A widely used approach in transformer models is to use embeddings of CLS tokens for sequence classification. We explored several ways to combine these vectors into a single representation:

- Concatenation of WT and MT embeddings of the mutation position (Fig. 1).
- Outer product of WT and MT embeddings of the mutation position (Fig. 2).
- Linear combination WT and MT embeddings of the mutation position (Fig. 3).
- Linear combination of CLS embeddings (Fig. 3).
- Linear combination of CLS embeddings concatenated with WT and MT embeddings of mutation position (Fig. 3).

**Fig. 2.**
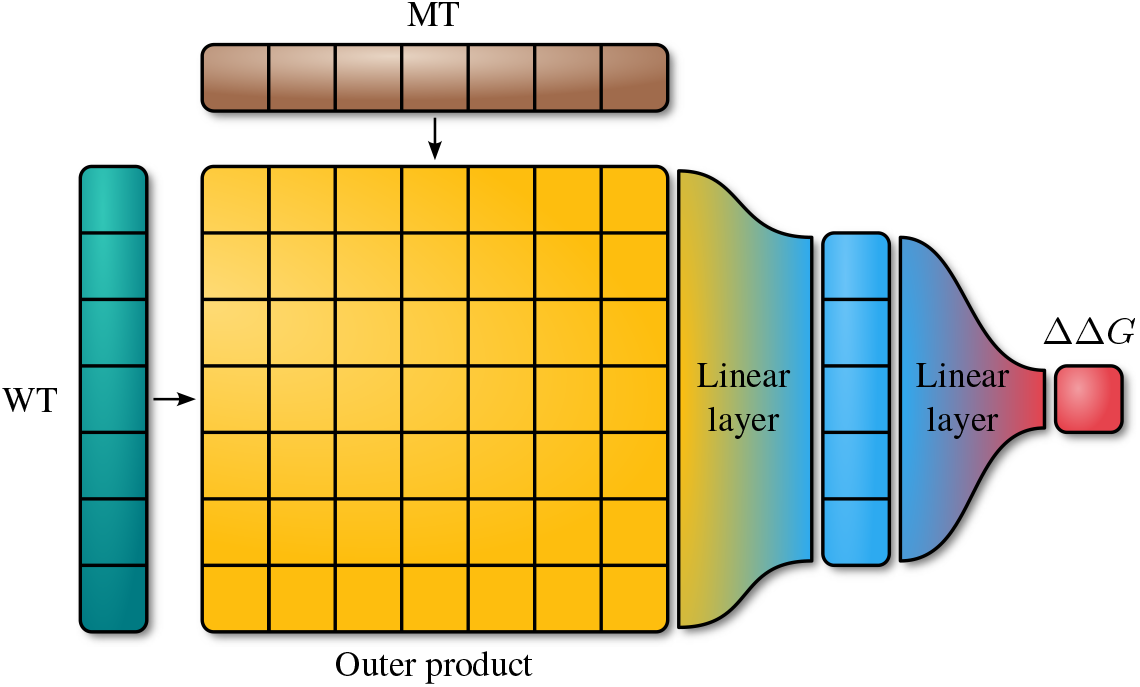
Architecture of the model that uses the outer product of token embeddings in mutation position of wild-type (WT) and mutated (MT) protein passed as input to the neural network with one hidden layer.

**Fig. 3.**
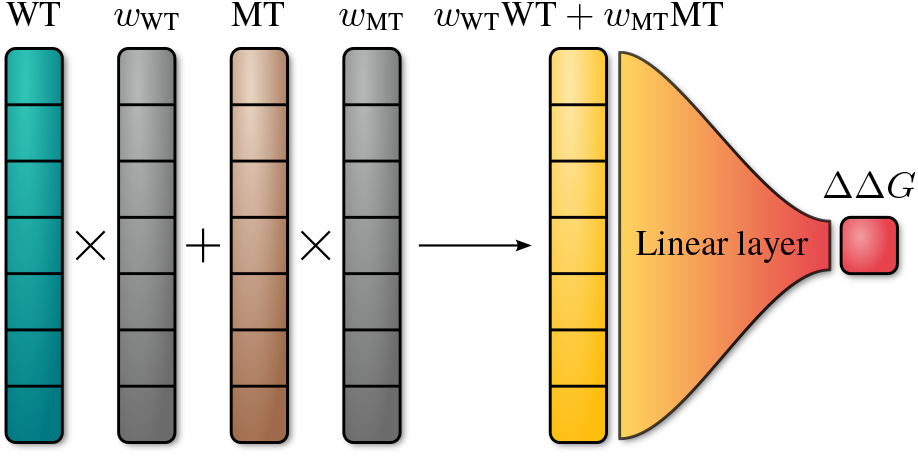
Architecture of the models that use the linear combination of embeddings of wild-type (WT) and mutated (MT) protein with vector weights *w*_WT_,_MT_. Multiplication of token embeddings with weight vectors is performed element-wise.

### 2.4 Model training and ensembling

All the models where trained for 3 epochs with the ADAM optimizer and a batch size of one. The learning rate was increased linearly from 0 to 1e-5 during the first 30% of the examples and then linearly decreased to 0 for the remaining examples. We did not freeze the transformer backbone and trained all model weights in an end-to-end manner.

To increase stability and improve the quality of predictions, we used an ensemble of all five models with different regression heads described previously. The final predictions were made by averaging the predictions of the five individual models in the ensemble.

### 2.5 Model evaluation

We used Pearson correlation (*r*), root mean square error (RMSE), and mean absolute error (MAE) to evaluate the performance of PROSTATA and compare it with previously developed methods. These metrics are used in original articles on other methods and in reviews. Therefore, to compare PROSTATA with other publicly available tools on original data sets, we used the performance metrics accordingly with the corresponding articles (Li et al., 2020; Benevenuta et al., 2021; Wang et al., 2021). Performance metrics of various models on Ssym and S669 datasets are taken from the work by Pancotti C. et al. (Pancotti et al., 2022).

## 3 Results and discussion

### 3.1 Regression head comparison

We compared the performance of different regression head architectures using 5-fold cross-validation. We used the protein cluster data to build the splits for cross-validation with each cluster assigned to a single fold. This ensured that for each fold, the examples in the test set were different from the ones in the training set. The results show that none of the models has a clear advantage over the others, while ensemble of five models have the highest performance (Table 1).

**Table 1.** Results of 5-fold cross-validation for different regression heads.

| Embedding | Merge | Pearson r | RMSE | MAE |
| --- | --- | --- | --- | --- |
| position | outer product | 0.65 | 1.65 | 1.12 |
| position | concatenation | 0.64 | 1.64 | 1.10 |
| CLS | linear | 0.66 | 1.70 | 1.44 |
| position | linear | 0.67 | 1.60 | 1.06 |
| CLS+position | linear | 0.67 | 1.60 | 1.06 |
| Ensemble |  | **0.69** | **1.57** | **1.03** |

### 3.2 Effects of asymmetrical datasets

We examined the effect of the regression head selection on how the model learns the symmetry effects from both symmetric and non-symmetric datasets. For this, we trained our models on Q3488 and Q3421 datasets and tested on Ssym. The Q3488 dataset contains an equal number of destabilizing and stabilizing mutations, while the Q3421 dataset is heavily biased toward destabilizing mutations. Moreover, Q3488 dataset does not contain proteins homologous to ones in the Ssym dataset. The results are presented in Table 2.

**Table 2.** Results of models trained on non-symmetric (Q3421) and symmetric (Q3488) sets and tested on the Ssym set.

| Embedding | Merge | Q3421 |  | Q3488 |  |
| --- | --- | --- | --- | --- | --- |
| | | $r_{dir}$ | $r_{rev}$ | $r_{dir}$ | $r_{rev}$ |
| position | outer product | 0.84 | -0.55 | 0.49 | 0.50 |
| position | concatenation | 0.72 | -0.63 | 0.38 | 0.43 |
| CLS | linear | 0.71 | 0.71 | 0.53 | 0.53 |
| position | linear | 0.89 | 0.89 | 0.43 | 0.43 |
| CLS+position | linear | 0.89 | 0.89 | 0.40 | 0.41 |
| Ensemble |  | 0.88 | 0.73 | 0.50 | 0.51 |

The models that use regression heads with linear merge of embeddings of wild-type and mutated sequences are able to learn the symmetry properly even when trained on a biased dataset. Models with outer product and concatenation merging are highly dependent on the balance in the training set and show negative correlation when presented with test set with bias reversed from training set. When provided with a balanced training set, all models are able to perform equally well on direct and reverse mutations.

For further analysis, we decided to use the ensemble of all 5 modes with different regression heads to ensure the ensemble diversity.

### 3.3 Comparison with peer neural network models

We compared the performance of our model with the results of other published neural network models. For this purpose we used training and test sets from the original articles. Among NN-based models we considered:

- ThermoNet predicts ΔΔ*G* using an ensemble of 3D CNN (Li et al., 2020). ThermoNet treats mutation site environments as multichannel voxel grids parameterized using atom biophysical properties.
- ACDC-NN-Seq is a convolutional neural network to predict changes in protein stability based solely on the protein sequence, unlike its predecessor ACDC-NN that uses 3D structural information (Benevenuta et al., 2021). ACDC-NN-Seq takes as input both direct and reverse variations, extracts features using convolution operations, and then feeds them into two differential siamese neural networks.
- ProS-GNN (Wang et al., 2021) is a deep graph neural network that was incorporated into BayeStab (Wang et al., 2022), a Bayesian neural network predicting ΔΔ*G* and evaluating uncertainty of its predictions.

The comparison on original datasets allows to assess the performance of different model architectures under comparable conditions. To compare the models we used Pearson correlation coefficient and RMSE metrics. Metrics for reviewed models were taken from original articles (Li et al., 2020; Benevenuta et al., 2021; Wang et al., 2021).

The results show that PROSTATA outperforms most neural network models on the Ssym, p53 and myoglobin datasets (Table 3). At the same time PROSTATA demonstrates comparable performance with ThermoNet. It has to be noted that ThermoNet was developed for structure-based prediction of effects of amino acid substitutions, whereas PROSTATA uses only sequence-based input.

**Table 3.** Neural network models on corresponding training and test sets.

| Train | Test | Pearson r |  | RMSE |  |
| --- | --- | --- | --- | --- | --- |
|  |  | Orig<sup>1</sup> | Ours | Orig<sup>1</sup> | Ours |
| *ThermoNet* |  |  |  |  |  |
| Q3488 | Ssym | 0.47 | **0.50** | 1.56 | **1.44** |
| Q3488 | Ssym<sub>r</sub> | 0.47 | **0.51** | 1.55 | **1.39** |
| *ACDC-NN* |  |  |  |  |  |
| S2648 + Vb | Ssym | 0.61 | **0.87** | 1.48 | **0.79** |
| S2648 + Vb | Ssym<sub>r</sub> | 0.61 | **0.77** | 1.48 | **1.43** |
| *ACDC-NN-Seq* |  |  |  |  |  |
| S2648 + Vb | Ssym | 0.55 | **0.87** | 1.44 | **0.79** |
| S2648 + Vb | Ssym<sub>r</sub> | 0.55 | **0.77** | 1.44 | **1.43** |
| S2648 + Vb | Myoglobin | 0.56 | **0.65** | 0.97 | **0.83** |
| S2648 + Vb | p53 | **0.67** | 0.60 | 1.62 | **1.50** |
| *ProS-GNN* |  |  |  |  |  |
| S2648 | Ssym | 0.61 | **0.83** | 1.23 | **0.94** |
| S2648 | Ssym<sub>r</sub> | 0.56 | **0.63** | **1.30** | 1.64 |
| S2648 | Myoglobin | 0.48 | **0.64** | 1.27 | **0.85** |
| S2648 | Myoglobin<sub>r</sub> | 0.43 | **0.64** | 1.19 | **1.05** |
| Q3421 | Ssym | 0.51 | **0.88** | 1.47 | **0.74** |
| Q3421 | Ssym<sub>r</sub> | 0.51 | **0.73** | **1.43** | 1.48 |
| Q3421 | Myoglobin | 0.51 | **0.70** | 1.20 | **0.85** |
| Q3421 | Myoglobin<sub>r</sub> | 0.45 | **0.48** | 1.31 | **1.10** |

<sup>1</sup>Metrics for reviewed models were taken from original articles (Li et al., 2020; Benevenuta et al., 2021; Wang et al., 2021).

Overall this data demonstrated that pretrained protein language models outperform most of other approaches based on neural networks.

However the large difference in PROSTATA performance metrics obtained in this analysis can be explained by different size as well as presence of homology of proteins in the train and test sets. In particular the largest correlation of 0.88 was obtained for S2648+Vb train set and Ssym test sets and the lowest one for Myoglobin reverse mutations. This highlights an importance of enhancing number of samples in the train set as well as excluding homologous proteins from the test set.

### 3.4 Evaluation on common test sets

Models that predict the effect of single mutations on protein stability are commonly benchmarked on Ssym dataset. The greatest challenge of these estimations is the overlap between the training set and the test set, leading to inflated performance metrics (Li et al., 2020). Some models like Thermonet and SCONES specifically craft their train sets to avoid such an intersection. A recent review of available tools to predict the effect of single mutations on protein thermostability (Pancotti et al., 2022) introduced a new S669 data set that contains proteins that are different from commonly used training sets. This data set allows for a fair comparison of different tools.

We evaluated our model on both S669 and Ssym datasets. From our dataset, we excluded proteins from train set with degree of identity higher than 75% compared to any test set protein. The results are presented in Table 4 and Supplementary Table S5, respectively. Pearson correlation coefficient of 0.48 was achieved by PROSTATA both for direct and reverse mutations in S669 dataset. To compare its performace with other tools we used metrics obtained from (Pancotti et al., 2022).Pearson correlation coefficient obtained by PROSTATA was higher compared to sequenced-based tools and comparable with metrics of structure-based tools.

**Table 4.** Performance of the models on the S669 dataset.

| Model | Direct |  |  | Reverse |  |  |
| --- | --- | --- | --- | --- | --- | --- |
|  | r | RMSE | MAE | r | RMSE | MAE |
| PROSTATA | 0.48 | 1.44 | 0.99 | 0.48 | 1.44 | 0.99 |
| INPS-Seq | 0.43 | 1.52 | 1.09 | 0.43 | 1.53 | 1.1 |
| ACDC-NN-Seq | 0.42 | 1.53 | 1.08 | 0.42 | 1.53 | 1.08 |
| DDGun | 0.41 | 1.72 | 1.25 | 0.38 | 1.75 | 1.25 |
| *ACDC-NN* | 0.46 | 1.49 | 1.05 | 0.45 | 1.5 | 1.06 |
| *DDGun3D* | 0.43 | 1.6 | 1.11 | 0.41 | 1.62 | 1.14 |
| *PremPS* | 0.41 | 1.5 | 1.08 | 0.42 | 1.49 | 1.05 |
| *ThermoNet* | 0.39 | 1.62 | 1.17 | 0.38 | 1.66 | 1.23 |
| *Rosetta* | 0.39 | 2.7 | 2.08 | 0.4 | 2.68 | 2.02 |
| *Dynamut* | 0.41 | 1.6 | 1.19 | 0.34 | 1.69 | 1.24 |
| *INPS3D* | 0.43 | 1.5 | 1.07 | 0.33 | 1.77 | 1.31 |
| *SDM* | 0.41 | 1.67 | 1.26 | 0.13 | 2.16 | 1.64 |
| *PoPMuSiC* | 0.41 | 1.51 | 1.09 | 0.24 | 2.09 | 1.64 |
| *MAESTRO* | 0.5 | 1.44 | 1.06 | 0.2 | 2.1 | 1.66 |
| *DUET* | 0.41 | 1.52 | 1.1 | 0.23 | 2.14 | 1.68 |

Metrics for reviewed models were taken from (Pancotti et al., 2022). Models in italic are structure-based.

The results on S669 show that our model improves by a large margin over existing solutions, both due to a new architecture and the use of a new dataset. Additionally, the PROSTATA model uses only the amino acid sequence as an input, without requiring explicit structural, evolutionary data, or additional features as some of the existing solutions.

### 3.5 Application

PROSTATA was developed to predict the effects of single point substitutions in proteins based on amino acid sequences alone. The accuracy of the model should depend primarily on the embeddings derived from the pre-trained protein language model. Protein language models are known to capture structural and evolutionary features (Hie et al., 2022; Lin et al., 2022) and thus PROSTATA is expected to be applicable for a variety of protein cases. To evaluate the applicability spectrum of PROSTATA, we measured its performance in a range of difficult cases.

In particular, we tested the predictive capacity of PROSTATA for mutants according to its location within the protein structure, the oligomerization state of protein, the solvent solubility, and the presence of small-molecule binding sites. In the first experiment, the mutant positions of the S669 test set were classified according to the location within the protein structure based on the solvent accessibility of the amino acid residues (Fig 4A-C) and corresponding secondary structure elements.

**Fig. 4.**
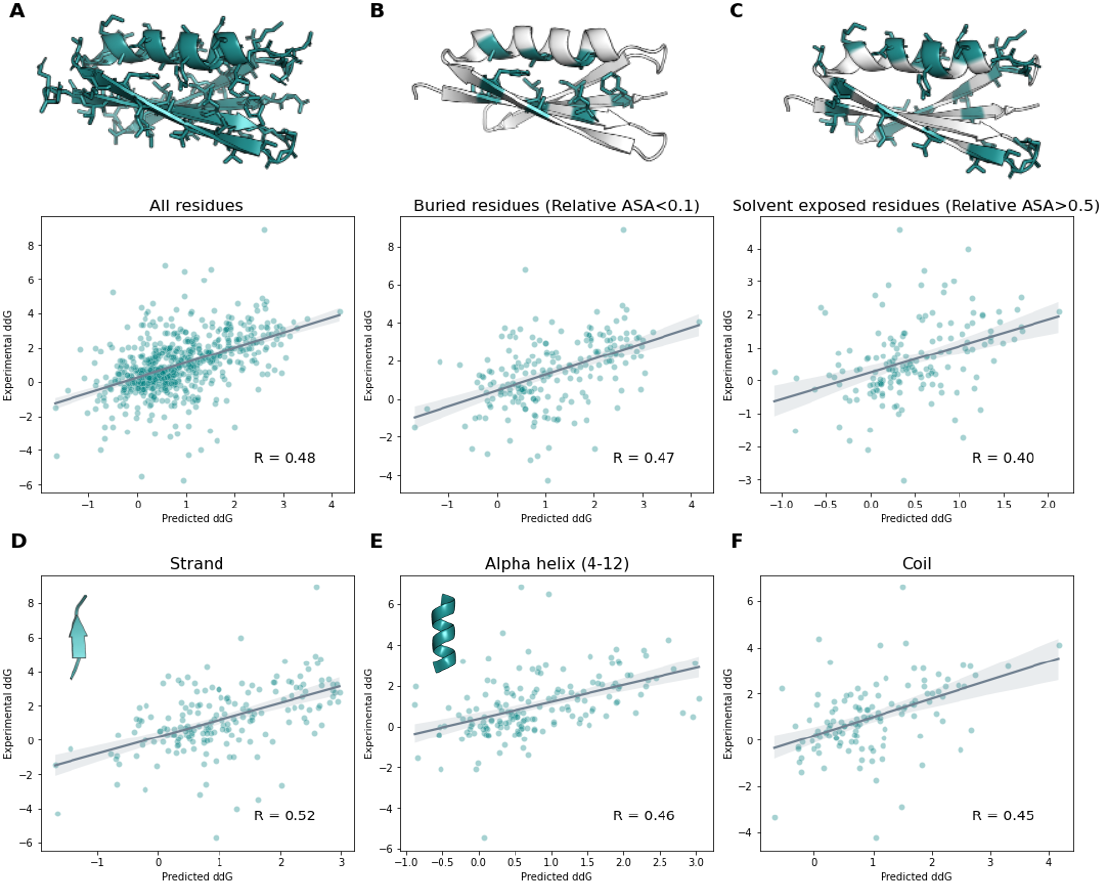
PROSTATA predictions on the S669 test set. (A-C) Comparison of PROSTATA performance for buried and solvent exposed mutant residues. Correlation between predicted and experimental ΔΔ*G* values for all residues (A), buried residues (B) and solvent exposed residues (C). Regions corresponding to the denoted condition are colored in cyan on the top. (D-F) Comparison of PROSTATA performance for mutants according to the corresponding element of the secondary structure. Correlation between predicted and experimental ΔΔ*G* values for Strand (D), Helix (E), Coil (F). The Pearson correlation coefficient is denoted on the bottom right corner. Representative structures are snow on the top. Relative ASA and secondary structure elements were predicted using DSSP tool (Kabsch and Sander, 1983).

We observed that the correlation of the experimental and predicted values was higher for mutant amino acid residues buried in the protein structure compared to that for solvent exposed residues. This dependence also manifests itself in low perplexity values, which may be attributed to the higher evolutionary stability of buried residues located in well-structured regions. Beta-strands and alpha-helices (4-12) are the most common secondary structure elements within the experimentally resolved structures. For beta-strand regions, PROSTATA demonstrated the best performance, with a slightly lower performance for alpha-helices and coils.

Several proteins included in the data set have a well-packed tertiary fold under biologically relevant conditions only in the oligomeric form. In particular, amyloid peptides are known to be disordered as monomers. Other proteins could be prone to form homodimers or other states of homoligomerization (Fig. 5A). To analyze the performance of PROSTATA on such cases, we developed a test set that includes oligomeric proteins. Proteins were considered oligomeric if they have at least 30% residues that interact with other subunits in the experimentally resolved structure within radii of 4.5 Å. Several representative entities of this test set are shown on Fig 5A.

**Fig. 5.**
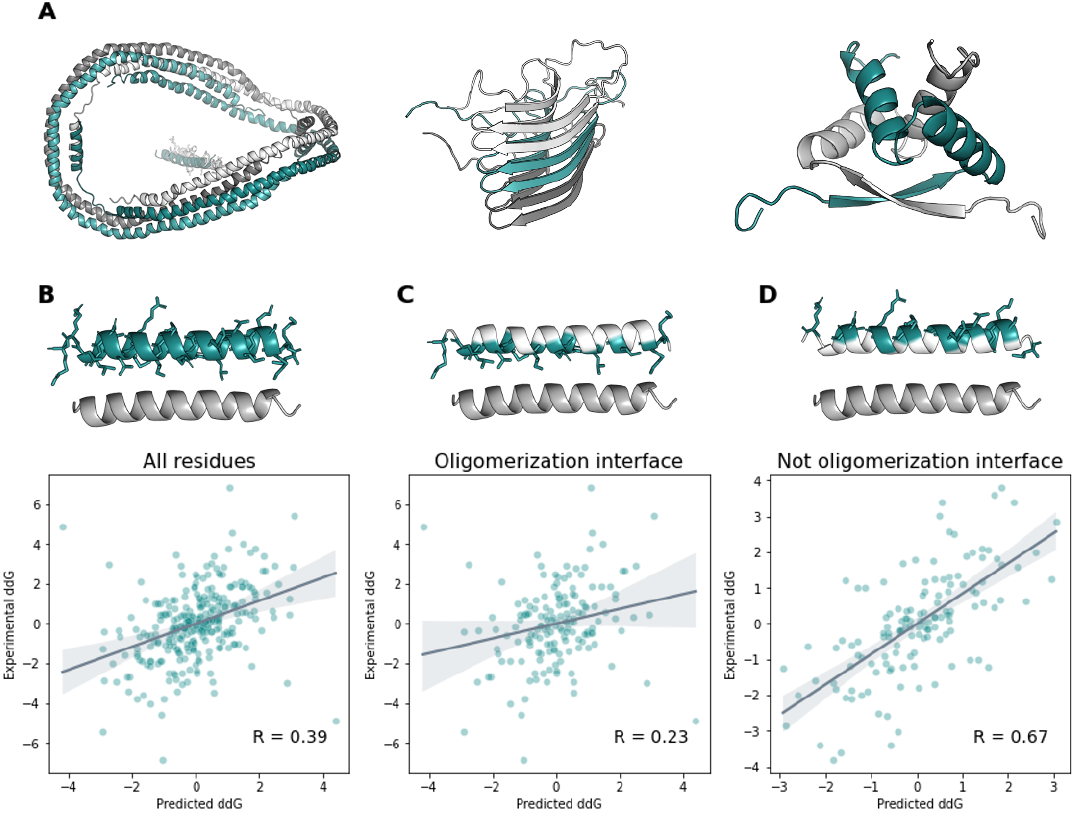
PROSTATA predictions for the test set of oligomeric proteins. (A) Representative examples of the test set, including homotrimer (left), amyloid (center), homodimer (right) structures. (B-D) Scatter plots for all mutant residues (B), mutant residues located on oligomerization protein-protein interaction interface (C), and not oligomerization protein-protein interaction interface (D) are shown. The Pearson correlation coefficient is denoted on the bottom right corner. Representative structures are snow on the top. Regions corresponding to the condition are colored dark green. Test set included following PDB codes: 1UWO_A, 1R6R_A, 2KJ3_A, 1SCE_A, 1SAK_A, 1ARR_A, 1ZNJ_A, 2A01_A, 2H61_A, 1CDC_B, 1BFM_A, 1ZNJ_B, 1AV1_A, 3MON_B.

As expected, the correlation between the experimental and predicted ΔΔ*G* values for such test set was lower compared to the original test set. Furthermore, PROSTATA showed low performance only in predicting changes in the ΔΔ*G* values for substitutions located at the protein-protein interaction interface. This is expected since we did not provide any information on protein oligomerization for the model. Therefore, PROSTATA is suitable for monomeric proteins, whereas for oligomeric proteins, an approach with explicit 3D structures may be more beneficial (Fig. 6).

**Fig. 6.**
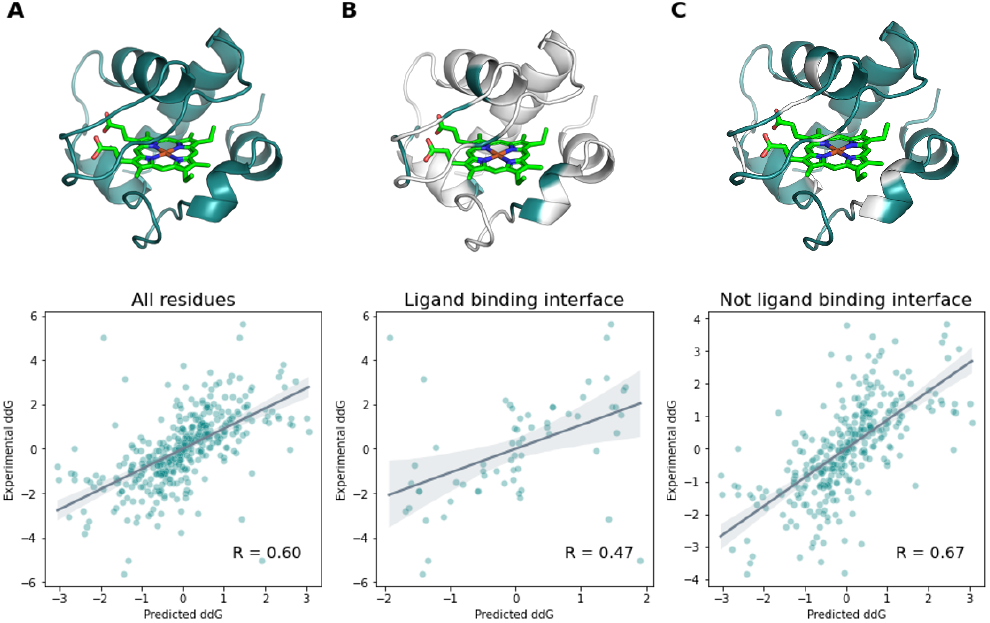
PROSTATA predictions for the class of proteins bound to hemoglobin or hemoglobin-derivatives. (A) Correlation between predicted and experimental ΔΔ*G* values for all residues, (B) ligand binding interface residues, (C) not ligand binding interface residues. The Pearson correlation coefficient is denoted on the bottom right corner. Representative structures are snow on the top. Regions corresponding to the denoted condition are colored cyan. Hemoglobin atoms are shown in light green. Test set included following PDB codes: 1C52_A, 1YCC_A, 1CYO_A, 1C2R_A, 1B5M_A, 1AKK_A, 1I5T_A, 1BVC_A, 1YEA_A, 1CYC_A, 451C_A, 1A7V_A.

Other challenging cases might include predicting the effect of mutations at small molecule and cofactor binding sites. Cofactor binding is known to generally stabilize the protein fold. Its location is not explicitly encoded in protein language models, but might be captured implicitly due to evolutionary traits.

To study the performance of PROSTATA, we split the data set on all hemoglobin or hemoglobin derivative binding proteins and other proteins as a test and training set, respectively. Surprisingly, PROSTATA demonstrated above average precision in this class of proteins, indicating that the protein language model is likely to understand the considered class of proteins well (Fig. 6). This may be due to large number of hemaglobin binding proteins in the UniProt database. At the same time, as expected, the overall precision for ligand binding residues was lower than that for other residues.

## 4 Conclusion

In this paper, we used the transfer learning approach to build a predictive model based on combinations of embeddings from the pre-trained protein language model ESM2. The model, PROSTATA, is an ensemble of five models with different regression heads.

To train our model, we have prepared a new dataset. To do this, we combined publicly available datasets, filtered and harmonised data to eliminate contradictory entries. Additionally, to prevent data leakage, we excluded from the training set all proteins that are more than 75% identical to those in the test set.

Compared to other models on their respective datasets, PROSTATA showed higher performance in terms of correlation and error. Comparing the performance of different models on the S669 dataset revealed that PROSTATA outperforms rivals due to both the new architecture and the use of the new training dataset.

Analysis of the performance of our new model on protein classes known to be challenging in ΔΔ*G* prediction suggest that PROSTATA acquired broad domain knowledge through transfer. Thus, it can be successfully applied to other classes of proteins besides those commonly found in protein stability datasets.

This work opens opportunities to develop new models to solve similar predictive problems. One option, for example, is to replace a pre-trained model with a more advanced one or combine several different pre-trained models. Combining models pre-trained on different data or using different modalities, such as protein sequences, multiple sequence alignments, or 3D structure data, can further broaden the scope and improve the accuracy of predictions.

## Author Contributions

D.U. and F.N. wrote the code for the models. D.U. carried out the model evaluation. N.V.I. performed the model application analysis. T.I.S. and M.S. cleaned the samples and constructed the dataset. D.U, P.V.S., N.V.I., T.I.S., and O.L.K designed the study. P.V.S., T.I.S., D.U, N.V.I., and F.N. prepared the manuscript.

## Competing interests

No competing interest is declared.

## Data Availability

The datasets generated for this study can be found in the GitHub repository: https://github.com/AIRI-Institute/PROSTATA/tree/main/DATASETS and Supplementarty tables.

## Notes

### Competing Interest Statement

The authors have declared no competing interest.

